# Role of the Fourth Transmembrane Segment in TRAAK Channel Mechanosensitivity

**DOI:** 10.1101/463034

**Authors:** Mingfeng Zhang, Fuqiang Yao, Chengfang Pan, Zhiqiang Yan

**Affiliations:** State Key Laboratory of Medical Neurobiology, Human Phenome Institute, Ministry of Education Key Laboratory of Contemporary Anthropology, Collaborative Innovation Center of Genetics and Development, Department of Physiology and Biophysics, School of Life Sciences, Fudan University, Shanghai 200438, China; Academy for Advanced Interdisciplinary Studies, Peking University, Beijing 100871, China

## Abstract

Mechanosensitive ion channels such as Piezo, TRAAK, TRPs and OSCA are important transmembrane proteins that are involved in many physiological processes such as touch, hearing and blood pressure regulation. Unlike ligand-gated channels or voltage-gated ion channels, which have a canonical ligand-binding domain or voltage-sensing domain, the mechanosensitive domain and related gating mechanism remain elusive. TRAAK channels are mechanosensitive channels that convert a physical mechanical force into a flow of potassium ions. The structures of TRAAK channels have been solved, however, the functional roles of the structural domains associated with channel mechanosensitivity remains unclear. Here, we generated a series of chimeric mutations between TRAAK and a non-mechanosensitive silent TWIK-1 K2P channel. We found that the selectivity filter region functions as the major gate of outward rectification and found that lower part of fourth transmembrane domain (M4) is necessary for TRAAK channel mechanosensitivity. We further demonstrated that upper part of M4 can modulate the mechanosensitivity of TRAAK channel. Furthermore, we found that hydrophilic substitutions of W262 and F121 facing each other, and hydrophobic substitutions of Q258 and G124, which are above and below W262 and F121, respectively, greatly increase mechanosensitivity, which suggests that dynamic interactions in the upper part of M4 and PH1 domain are involved in TRAAK channel mechanosensitivity. Interestingly, these gain-of-function mutations are sensitive to cell-poking stimuli, indicating that cell-poking stimuli generate a low membrane mechanical force that opens TRAAK channels. Our results thus showed that fourth transmembrane domain of TRAAK is critical for the gating of TRAAK by mechanical force and suggested that multiple dynamic interactions in the upper part of M4 and PH1 domain are involved in this process.

## Introduction

K2P channels are indispensable for background leak currents that regulate the membrane potential and excitability of many different cell types. The channel activities are regulated by many physiological factors, such as pH, lipids, temperature, and mechanical force [1]. Fifteen K2P homologues exist in humans and can be structurally and functionally divided into six subfamilies, namely, the TWIK, TREK, TASK, THIK, TALK, and TRESK subfamilies. Among these six subfamilies, three members of the TREK subfamily, the TREK-1, TREK-2, and TRAAK channels, are activated by stimuli from a physical mechanical force [2]. The results of experiments involving TREK channels functionally reconstituted in liposomes together with the lysophospholipids that activate TREK channels indicate that TREK channels sense the physical mechanical force in the lipid membrane environment [3, 4]. Recently, the crystal structures of the human TRAAK channel suggest a lipid seal mechanism for the mechanical force sensing of TRAAK channels [5]. The crystal structures of TREK-2 also reveal that lipids in the channel cavity play an important role in channel gating, which is consistent with the lipid seal mechanism [6].

In this study, we attempted to identify the critical elements in TRAAK channel mechanosensitivity by generating a series of chimeric mutations between mechanosensitive TRAAK and a non-mechanosensitive silent TWIK-1 channel. Replacing the pore helix and selectivity filter (PH2-filter), the fourth transmembrane helix (M4) of TRAAK or only M4 of TRAAK with the corresponding residues of TWIK-1 revealed that M4 is indispensable for TRAAK channel mechanosensitivity and that the selectivity filter region is the major gate for outward rectification. Moreover, the results by replacing upper or lower part M4 of TRAAK with corresponding residues of TWIK-1 are consistent with the idea that the lower part of M4 is involved in sensing the mechanical force. Additionally, we found that hydrophilic substitution of W262 and F121, which are facing each other, and hydrophobic substitution of Q258 and G124, which are above and below W262 and F121, respectively, largely increase channel mechanosensitivity, which suggests that dynamic interactions in the upper part of M4 and PH1 domain are involved in mechanical gating of TRAAK channel. Interestingly, the channels with these gain-of-function mutations are more sensitive to cell-poking stimuli than wild-type TRAAK channels, indicating that cell-poking stimuli generate a low membrane mechanical force that opens TRAAK channels.

### Chimeras reveal that M4 is indispensable for TRAAK channel mechanosensitivity and that the selectivity filter region is the major gate for outward rectification

The fifteen K2P homologues display different functions despite their structural and sequence conservation [1]. Structural and sequence conservation provides the possibility of generating chimeric mutations between the different K2P channels. To explore the structural basis of channel mechanosensitivity, we first divided the K2P channel into eight segments by structural information: the first transmembrane helix (M1), the two-cap domain (Cap), the first pore helix and selectivity filter (PH1-filter), the second transmembrane helix (M2), the third transmembrane helix (M3), the second pore helix and selectivity filter (PH2-filter), the fourth transmembrane helix (M4) and the C-terminal domain (C-ter) (S1 Fig). We generated the chimeric mutations with a previously reported non-mechanosensitive silent TWIK-1 K2P channel [7, 8] and measured the background whole cell currents and negative pressure activated microscopic currents to study the effects of the eight segments on channel activity. Most chimeric mutations showed silent channel characteristics similar to TWIK-1 channels, and some mutations were similar to wild-type TRAAK channels (S2 Fig). Interestingly, replacing the PH2-filter and M4 of TRAAK with the corresponding residues of TWIK-1 (2-5 mutation) eliminated both the mechanosensitivity and outward rectification properties, resulting in large linear background potassium currents compared to those of wild-type TRAAK channels (Fig 1). Only replacing M4 of TRAAK with the corresponding residues of the TWIK-1 channel (2-6 mutation) eliminated mechanosensitivity but retained the outward rectification properties, with large voltage-dependent background potassium currents compared to those of wild-type TRAAK channels (Fig 1). The two non-mechanosensitive chimeras, the 2-5 and 2-6 mutations, suggested that M4 is critical for channel mechanosensitivity. Additionally, comparison of the 2-5 and 2-6 mutations clearly showed that the PH2-filter plays an important role in outward rectification. These results strongly suggest that the selectivity filter region functions as the major gate for outward rectification and that M4 of TRAAK might function as the mechanical force sensing region, which supports the C-type gate activation model [9-11].

**Fig 1.**
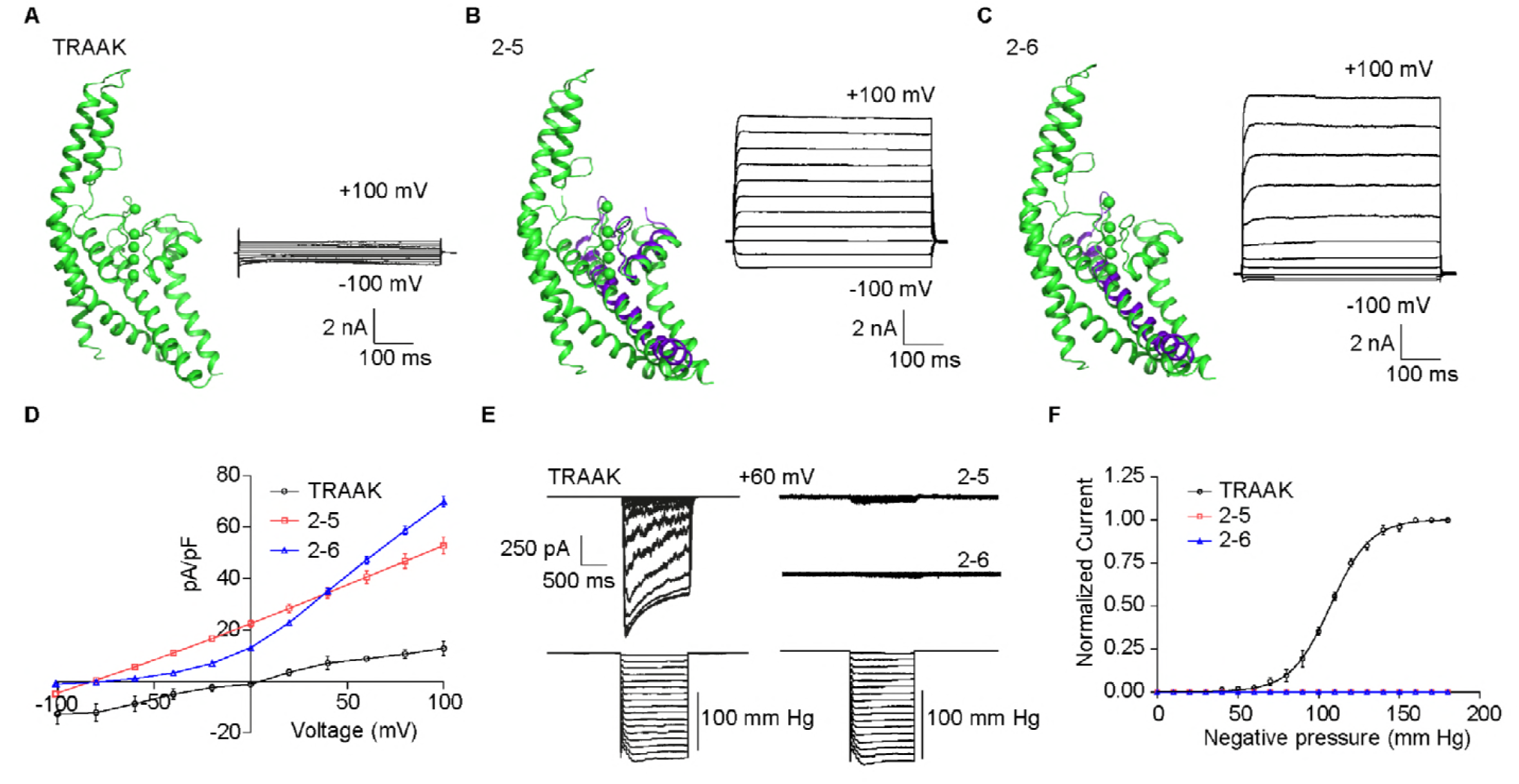
Chimeras reveal that M4 is indispensable for TRAAK channel mechanosensitivity, and the selectivity filter region is the major gate for outward rectification. (A-C) Schematic of a single subunit of the wild-type human TRAAK channel (A), 2-5 (B) and 2-6 (C) mutations (left). Violet segments are the corresponding residues of the TWIK-1 channel. Representative traces of wild-type human TRAAK channel (A) and the channels with the 2-5 (B) and 2-6 mutations (C) with (right) whole cell currents from -100 mV to +100 mV with +20 mV per step in HEK293T cells. (D) Statistical analysis of the current density of wild-type human TRAAK channel (black) and the channels with the 2-5 (red) and 2-6 mutations (blue), as well as the voltage relationships from A-C. Note the outward rectified behavior of the channel with the 2-6 mutation and the ohmic behavior of the channel with the 2-5 mutation. (E) Representative traces of negative pressure-activated wild-type human TRAAK (left) and 2-5 and 2-6 mutation (right) currents at the membrane potential of +60 mV. Negative pressure was applied from 0 mm Hg to the negative pressure until the currents reaching the maximum state with steps of 10 mm Hg. No dominant negative pressure-activated currents were detected with the 2-5 and 2-6 mutations. (F) Negative pressure-activated currents of the wild-type human TRAAK and the 2-5 and 2-6 mutations at the indicated membrane potential were fitted with the Boltzmann equation. P_50_ is 101.0 ± 5.4 mm Hg for the wild-type human TRAAK channel.

### The lower part of M4 is necessary for in mechanical gating and the upper part modulates the mechanosensitive currents of TRAAK

To further study the functional role of M4 in channel mechanosensitivity, we divided M4 into the upper part and the lower part (Fig 2 and S2 Fig). Interestingly, replacing the lower part of the M4 region of TRAAK with the corresponding residues of the TWIK-1 channel (2-6-2 mutation) eliminated the mechanosensitivity and resulted in silent properties (Fig 2A, 2D and 2E), indicating that the lower part of M4 is involved in mechanogating of the channel. Replacing the upper part of the M4 region with the corresponding residues of the TWIK-1 channel (2-6-1 mutation) reduced the half activation pressure threshold (P_50_) from 101.0 ± 5.4 mm Hg to 38.8 ± 3.8 mm Hg (Fig 2E and 2F), indicating an increase in channel mechanosensitivity. The 2-6-1 mutation also resulted in large voltage-dependent background potassium currents compared to those of the wild-type TRAAK channel (Fig 2B and 2D), suggesting that the mutation breaks some energy barrier for mechanotransduction. We also replaced the M4 upper part of the TREK-1 channel with the corresponding sequences of the TWIK-1 channel, and we found that the chimera had properties similar to the 2-6-1 mutation, which showed increased background potassium channel currents (Fig 2C and 2D). These results suggest that the lower part of M4 might be the core force-sensing domain of TRAAK, and the upper part of M4 modulates the channel activity.

**Fig 2.**
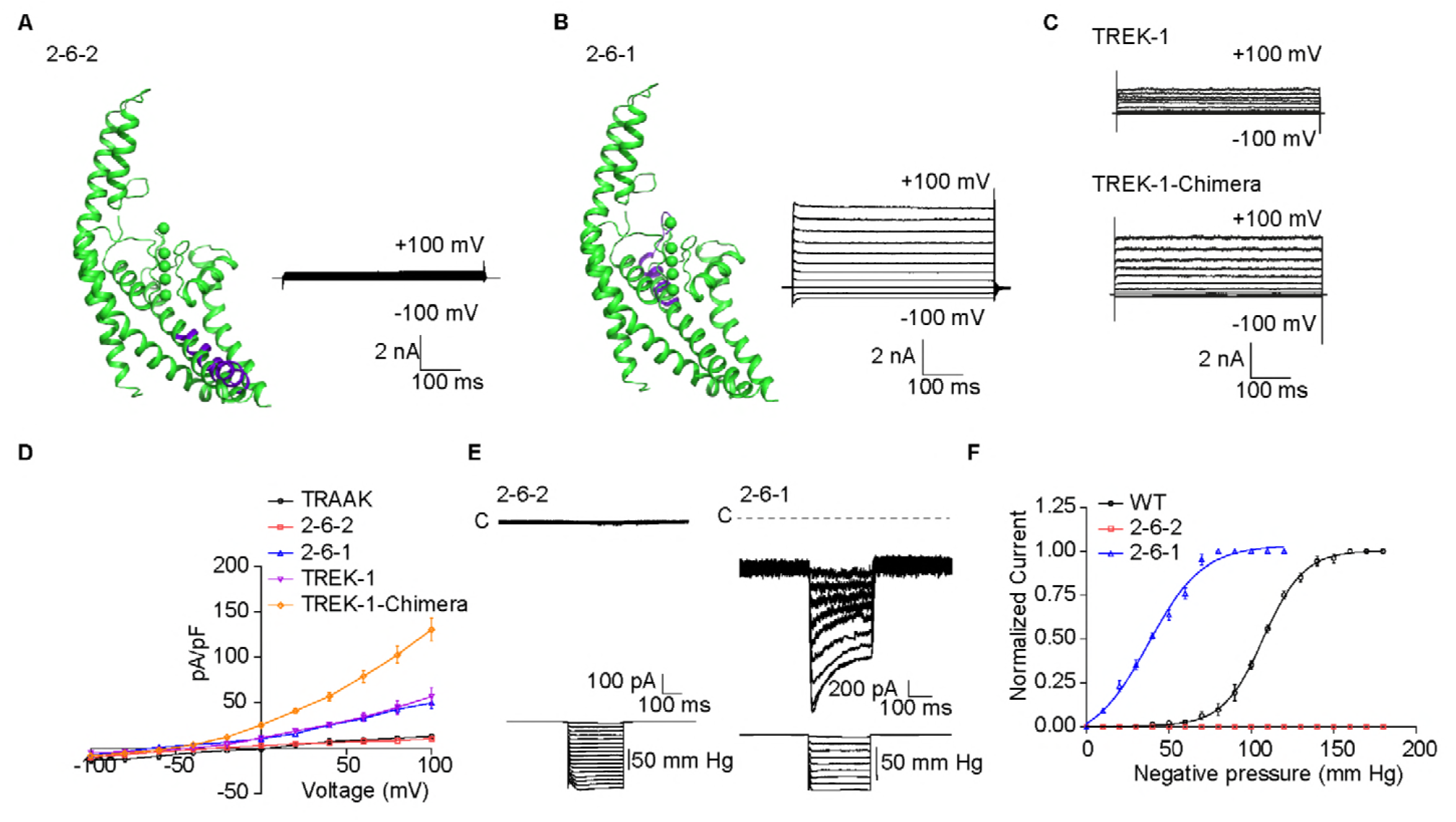
The lower part of M4 is necessary for in mechanical gating and the upper part of M4 modulate the mechanosensitive currents of TRAAK. (A-B) Schematic of a single subunit for the 2-6-2 (A) and 2-6-1 (B) mutations (left). Representative traces of 2-6-2 (A) and 2-6-1 (B) mutation (right) whole cell currents from -100 mV to +100 mV with +20 mV steps in HEK293T cells. (C) Representative traces of wild-type human TREK-1 (upper) and TREK-1 (upper part of M4 = TWIK-1) chimera mutation (lower) whole cell currents from -100 mV to +100 mV with +20 mV steps in HEK293T cells. (D) Statistical analysis of the current density of the wild-type human TRAAK (black), 2-6-2 (red), 2-6-1 (blue), wild-type human TREK-1 (violet) and TREK-1 (upper part of M4 = TWIK-1) chimera (yellow) mutation channels and the voltage relationships from A-C. (E) Representative traces of negative pressure-activated 2-6-2 (left) and 2-6-1 mutation (right) currents at the membrane potential of +60 mV. Negative pressure was applied from 0 mm Hg until the currents reached the maximum state with steps of 10 mm Hg. No dominant negative pressure-activated currents were detected for the 2-6-2 mutation. (F) The negative pressure-activated currents of wild-type human TRAAK, 2-6-2 (red) and 2-6-1 (blue) mutation channels at the indicated membrane potential were fitted with the Boltzmann equation. P_50_ is 101.0 ± 5.4 mm Hg for the wild-type human TRAAK and 38.8 ± 3.8 mm Hg for the 2-6-1 chimeric mutation.

To study the action of the upper part of M4 in regulating channel activity at the single residue level, we mutated residues in the M4 upper part the TRAAK channel into the corresponding residues of the TWIK-1 channel. The nonconserved sequences of the upper part of M4 comprise a small flexible linker (248-256) and an alpha helix (258-265) (Fig 3A and S2 Fig). We found that both the flexible linker (248-256) and the alpha helix (258-265) were involved in channel activity (Fig 3B, 3C and 3F).

**Fig 3.**
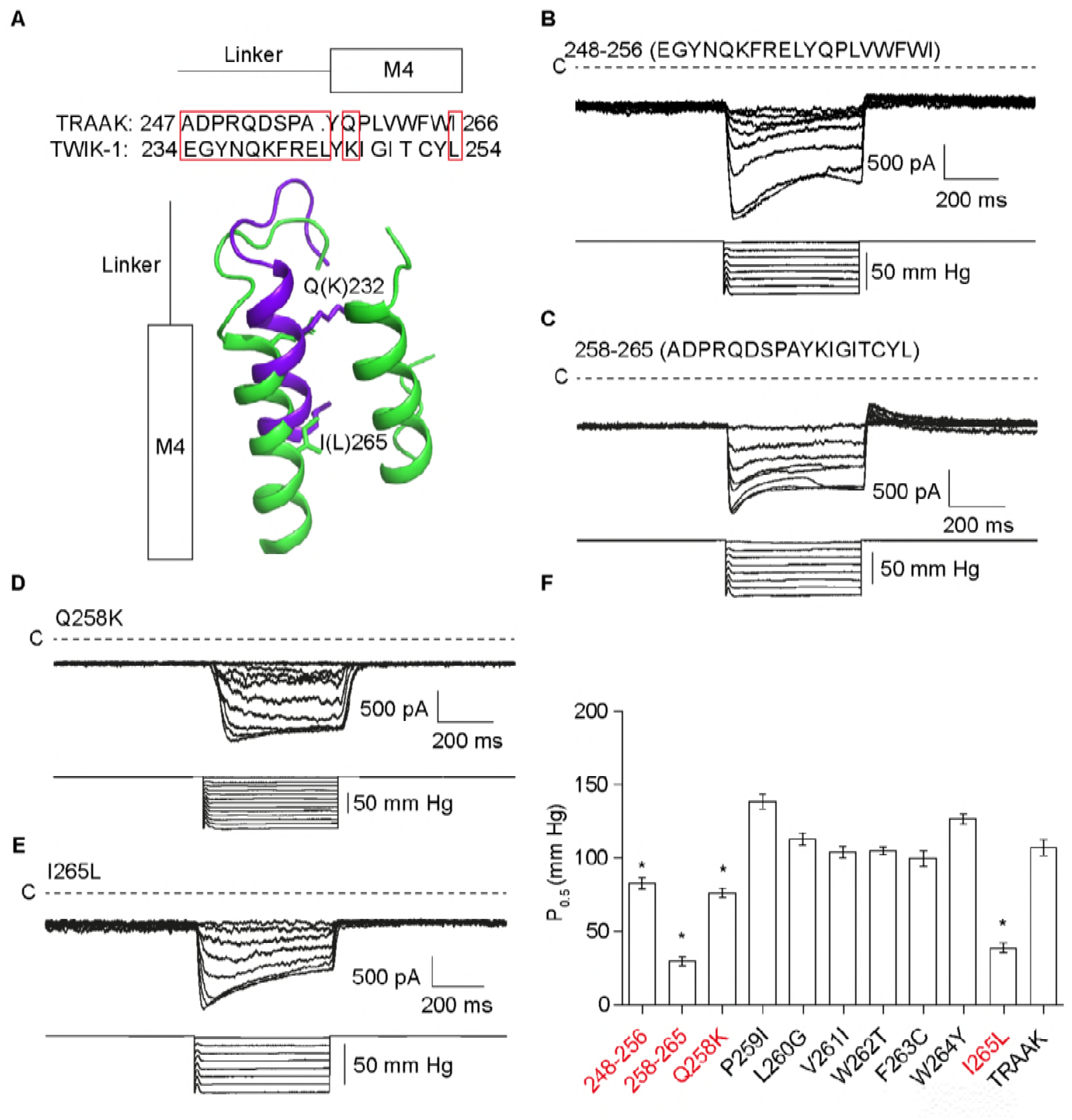
Multiple elements in the upper part of M4 regulate TRAAK channel activity. (A) Schematic in the upper part of M4. The gain-of-function (GOF) chimera mutations are indicated by red boxes. (B-E) Representative traces of negative pressure-activated 248-256, 258-265, Q258K and I265L mutation currents at the membrane potential of +60 mV. Negative pressure was applied from 0 mm Hg until the currents reached the maximum state, with steps of 10 mm Hg. (F) Statistical analysis of the P_50_ values of indicated mutations at the membrane potential of +60 mV. Statistical significance compared to wild type was calculated using Student’s t-test (*p < 0.05).

We next mutated residues in the alpha helix (258-265) of the TRAAK channel into the corresponding residues of the TWIK-1 channel step by step and found that Q258K and I265L both increased background potassium currents and mechanosensitivity (Fig. 3D, 3E and 3F). These results suggest that multiple elements in the upper part of M4 regulate channel activity.

### Hydrophilic substitution of W262 and F121 and hydrophobic substitution of Q258 and G124 largely increase channel mechanosensitivity

Previously, it was reported that W262, which is in the line of Q258 and I265, plays a critical role in TRAAK channel activities [12-14]. The I265L mutation produces a small side chain difference between isoleucine and leucine, while the gain-of-function mutations W262S and Q258K produce relatively different side chain properties. To identify the possible mechanisms of W262 and Q258 in channel activity, we mutated W262 and Q258 in the TRAAK channel into the 19 other amino acids. Mutations at W262 led to channels with varying P_50_ values that correlated with the hydrophobic value of substitutions (R^2^ = 0.5715, p < 0.0001) (Fig 4A). Substitutions of hydrophilic amino acids (W262D, W262E, W262N and W262Q) other than the positively charged amino acids (W262R and W262K) reduced the P_50_ value, while substitutions of hydrophobic amino acids (W262L, W262V and W262I) led to an increase in P_50_ (Fig 4A). Mutations at Q258 showed the opposite property as those at W262. The P_50_ value of Q258 mutations had a negative correlation with the hydrophobic value of substitutions of Q258 (R^2^ = 0.7035, p < 0.0001) (Fig 4B). Substitutions of hydrophobic amino acids (Q258W, Q258L, Q258V, Q258F and Q258I) and the positively charged amino acids (Q258R and Q258K) mainly reduced the P_50_ value (Fig 4B).

**Fig 4.**
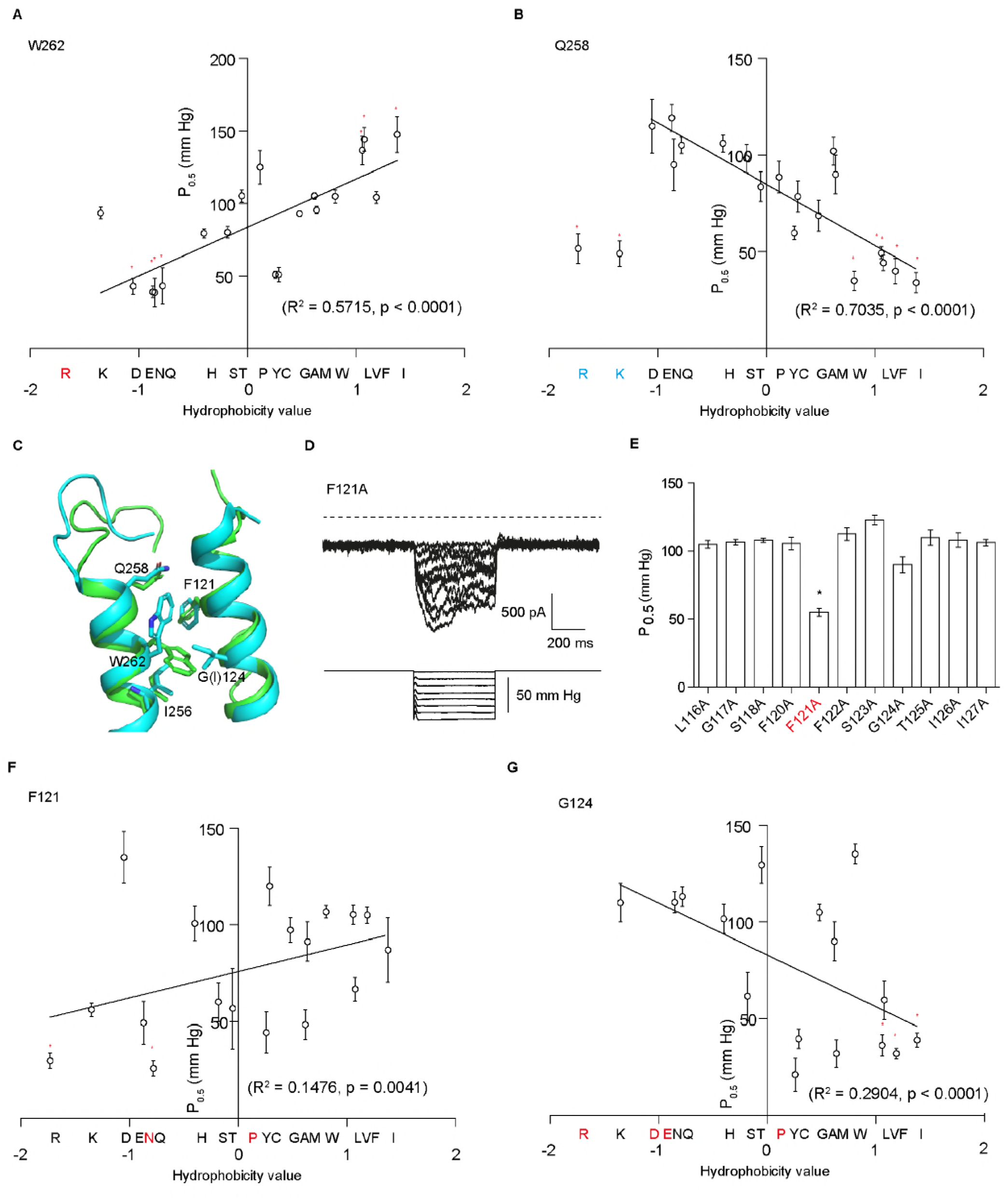
Hydrophilic substitutions of W262 and F121 and hydrophobic substitutions of Q258 and G124 largely increase the channel mechanosensitivity. (A-B) Statistical analysis of the relationships between P_50_ and hydrophobicity for the indicated W262 (A) and Q258 (B) substitution mutations. W262R (red) does not show channel activity. The Q258R and Q258K (blue) were removed for the curve fitting. There is a positive correlation between P_50_ and the hydrophobicity value for W262 (R^2^ = 0.5715, p < 0.0001) and a negative correlation for Q258 (R^2^ = 0.7035, p < 0.0001). Statistical significance compared to wild type was calculated using Student’s t-test (*p < 0.05). (C) Structural alignment of wild-type TRAAK (PDB ID: 3UM7) and G124I mutation (PDB ID: 4RUE) in the upper of M4 and PH1 domain. W262 and F121 show slight conformational changes in the upper of M4 and PH1 domain. (D) Representative traces of negative pressure-activated F121A mutation currents at the membrane potential of +60 mV. Negative pressure was applied from 0 mm Hg until the currents reached the maximum state, with steps of 10 mm Hg. (E) Statistical analysis of the P_50_ values of the indicated mutations at the membrane potential of +60 mV. Statistical significance compared to wild type was calculated using Student’s t-test (*p < 0.05). (F-G) Statistical analysis of the relationship between the P_50_ and hydrophobicity values for the F121 (F) and G124 (G) substitution mutations. F121N, F121P, G124R, G124D, G124E and G124P (red) do not show channel activity. There is a positive correlation between P_50_ and the hydrophobicity value for F121 (R^2^ = 0.1476, p = 0.0041) and a negative correlation for G124 (R^2^ = 0.2904, p < 0.0001). Statistical significance compared to wild type was calculated using Student’s t-test (*p < 0.05).

Previous studies suggested that W262 at M4 and G124 at PH1 may interact with each other [12, 13]. Q258 and W262 are in the upper part of M4, while Q258 is far away from G124 (Fig 4C). If there are multiple dynamic interactions in the upper part of M4 and PH1 domain, other critical sites may exist in this region. To determine the new critical in the PH1 domain, we performed an alanine scan of the PH1 domain. Interestingly, one alanine substitution mutation (F121A) led to a large increase in channel mechanosensitivity (Fig 4D and 4E). We also mutated F121 and G124 in the TRAAK channel to the 19 other amino acids. Interestingly, for F121, the relationship between its hydrophobicity value and P_50_ was similar to that of W262 (R^2^ = 0.1476, p = 0.0041) (Fig 4F), while G124 had a relationship similar to that of Q258 (R^2^ = 0.2904, p < 0.0001) (Fig 4G). The hydrophilic amino acid substitutions of F121 (F121R and F121Q) largely reduced P_50_ (Fig 4F), while the hydrophobic amino acid substitutions of G124 (G124L, G124F and G124I) reduced the P_50_ (Fig 4G). These results suggest that hydrophilic substitution of W262 and F121 and hydrophobic substitution of Q258 and G124 largely increase channel mechanosensitivity, indicating that multiple dynamic interactions in the upper part of M4 and PH1 domain are important in the gating of TRAAK channels.

### The GOF mutations in the upper part of M4 are sensitive to cell-poking stimuli

The cell-poking assay imitates touch behavior by using a glass tip to touch the cell, which converts mechanical force into the flow of ions. This assay is popular and successful in studying NompC and Piezo channels [15]. It is difficult to activate the TRAAK channel with a short displacement distance, while using cell-poking stimuli with a long displacement distance will disrupt the whole cell configuration [4]. In the Piezo channel, the P_50_ threshold is approximately -30 mm Hg, and the half activation displacement distance (D_50_) is approximately 5 μm [15, 16]. In our system, the results are consistent with the previously reported results of Piezo channels (Fig 5A). In the wild-type TRAAK channel, the P_50_ threshold is approximately 100 mm Hg (Fig 1F) approximately 3-4 times that of the *m*Piezo channel, and the D_50_ is difficult to determine despite the detection of visible cell-poking activated currents at long displacement distances (Fig 5B). In the gain-of-function mutations, the P_50_ value is approximately -30 mm Hg (Fig 4A, 4B, 4F and 4G), which is similar to the P_50_ threshold of the Piezo channel. We attempted to record the whole cell-poking activated currents of the representative gain-of-function mutations of 2-6-1, F121Q, Q258F and W236E (Fig 5). Comparison to the wild-type TRAAK channels revealed that the gain-of-function mutations make the channels sensitive to cell-poking stimuli (Fig 5), indicating that cell-poking stimuli and negative pressure shares a common mechanism to activate mechanosensitive channels, and cell poking has similar effect of actuating mechanosensitive channels with low negative pressure.

**Fig5.**
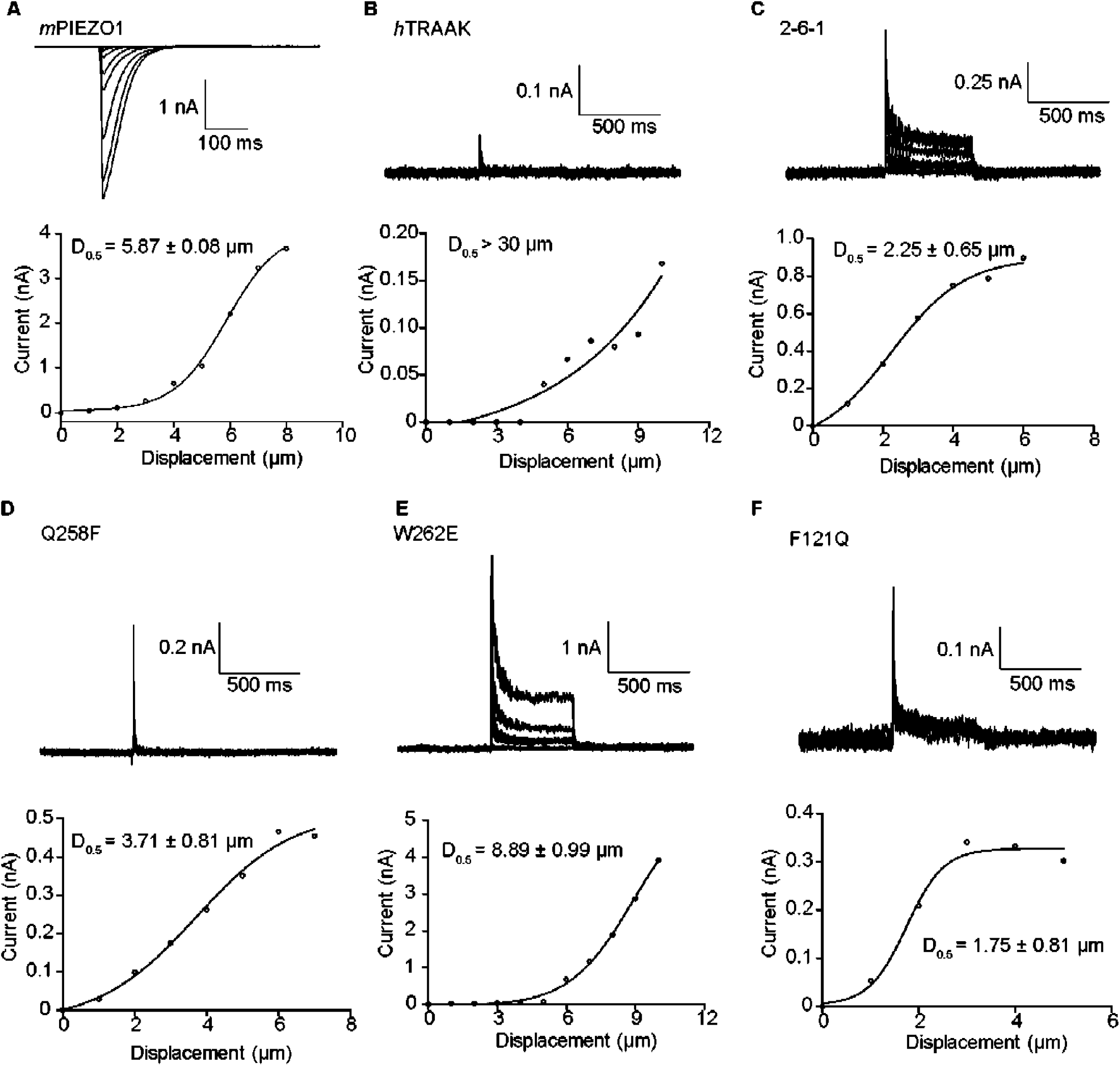
The GOF mutation channels in the upper of M4 and PH1 domain are sensitive to cell-poking stimuli. (A-F) Representative traces (upper) and statistical results (lower) of the currents from the cell poking-activated mPIEZO1 channel at the membrane potential of -60 mV and wild-type human TRAAK, 2-6-1, Q258F, W262E and F121Q mutation channels at the membrane potential of +60 mV. Cells were subjected to a series of mechanical steps via 1 μm movements of a stimulation pipette in the whole-cell patch mode. All recordings are representative of at least three separate experiments.

## Discussion

Mechanosensitive channels are widely distributed from prokaryotes to eukaryotes and are involved in the responses of many important physiological processes, such as touch hearing and blood pressure regulation [17]. In eukaryotes, only members of the TREK channel subfamilies are mechanosensitive potassium channels negatively regulating the cell excitability [1]. Reported stretch-activated mechanosensitive channels include OSCAs [18], Piezos [15], K2Ps [3, 19, 20], TRPs [21, 22], MscLs [23], and MscSs [24]. OSCA channels are dimer transporter-like family proteins. The M0 and M6 transmembrane helices of OSCA channels may play critical roles in channel mechanosensitivity [18]. Piezo channels are tetramers with long transmembrane helical units (THUs) assembled into a highly curved blade-like structure, and THUs may be involved in Piezo channel mechanosensitivity [25, 26]. K2P and TRP channels share a similar tetramer topology (one subunit comprises two pore helices and two selectivity filter p-loops in K2P) [5, 27]. The M4 transmembrane helix of K2P channels may serve as the force-sensing domain [5] (Fig 2). MscL channels are pentamers [28]. An iris-like expansion of MscL may be the channel gating mechanism [29]. MscS channels are heptamers [30]. Substantial rotational rearrangement of the three transmembrane helices occurs during channel activation [31]. These proteins, varying from dimer to heptamer, have no structural similarities and no recognizable force-sensing domain comparable to the voltage sensor or ligand-binding sites.

However, the electrophysiological properties of OSCA channels show some similarities to TREK channels [18]. Both have outward rectification properties, showing a “valve-like” channel behavior. Additionally, both show large-stretch activated currents and have similar relatively large P_50_ thresholds compared to that of the well-known mechanosensitive Piezo channel. Mechanosensitive OSCA channels share structural similarity with TMEM16A channels, which are calcium-activated chloride channels [18]. In TMEM16A channels, calcium induces the lower part of the sixth transmembrane helix (M6) to straighten, thus activating the channel [32]. OSCA channels may share a similar mechanism of straightening of the lower M6 region for channel activation [18]. M6 in OSCA and TMEM16A is also reminiscent of M4 in the TRAAK channel, hinting that the upper part of M6 may share functions similar to those of the upper part of M4.

C-type gating is a common means to respond to different physical inputs by changing the ion occupancy of the selectivity filter [9-11]. Large conformational changes of transmembrane helices are not necessary for C-type gating [10]. In K2P channels, slight conformational changes of the cryptic K2P modulator pocket converge at the selectivity filter C-type gate, which strongly affects the channel activity [33], which are consistent our results (Fig. 4C). At this point, when the structure is not available at the super-resolution level, functional studies of the channel are most instructive. The outward rectification property of the OSCA and TMEM16A channels and the external ions largely affect the channel activity phenotype of TMEM16A [34], similar with TRAAK channel, reminiscent of the “C-type gate” that may exist in OSCA and TMEM16A channels.

## Materials and methods

### Molecular biology

All channel DNA was used in the pIRES2eGFP vector. Human TRAAK and human TREK-1 DNA were obtained from GeneCopoeia, and the sequence was confirmed by Sanger sequencing. All mutations were generated by PCR and a DNA in-fusion protocol. HEK293T cells were maintained in DMEM with 5% FBS on poly-L-lysine-coated glass cover slips in 12-well plates. Cells were cotransfected using Lipofectamine 3000 (Invitrogen) with a total of 3 μg of DNA per 18-mm-diameter cover slip.

### Electrophysiology

HEK293T cells were transfected with plasmids and incubated for 24-36 h before recording. HEK293T cell electrophysiology measurements were performed in the indicated solution. The pipette resistance was 8-10 MΩ. For the cell-attached experiments, the bath solution contained 150 mM KCl, 5 mM EGTA and 10 mM Hepes (pH 7.2), and the pipette solution contained 142.5 mM NaCl, 7.5 mM KCl, 5 mM EGTA and 10 mM Hepes (pH 7.2). For the whole cell experiment, the bath solution contained150 mM NaCl, 5 mM EGTA and 10 mM Hepes (pH 7.2), and the pipette solution contained 142.5 mM KCl, 7.5 mM NaCl, 5 mM EGTA and 10 mM Hepes (pH 7.2). Recordings of patch-clamped cells were obtained using an Axopatch 700B (Molecular Devices) amplifier (Molecular Devices), filtered at 1 kHz and digitized at 10-100 kHz (Digidata 1440A, Molecular Devices). Negative pressure in the pipette was applied using a Suction Control Pro unit (Nanion) with a stepwise protocol through Clampex software. We used the I/Imax versus negative pressure curve approach to evaluate mechanosensitivity. Imax is the maximum current measured from the excised patch with cell-attached patch clamp settings. Glass probes for the whole-cell poking assay were made from borosilicate glass pipettes that were fire polished until becoming sealed with a tip diameter of approximately 3–4 μm. The probe was mounted to a piezo-driven actuator driven by a controller/amplifier controlled through Clampex software. After the formation of a whole-cell seal, the probe was positioned at 45° to the cell and gently attached to the cell membrane. All recordings were performed at a room temperature of 22°C. Data were then analyzed by pClamp10.4 software. All data were acquired from at least five independent cells.

## Acknowledgments

We thank all the Yan lab members for support. The research was supported by funds from the National Key R&D Program of China Project (Project 2017YFA0103900, 2016YFA0502800) to Z.Y., the Program for Professor of Special Appointment (Eastern Scholar of Shanghai, TP2014008) and the Shanghai Rising-Star Program (14QA1400800) to Z.Y. The National Natural Science Foundation of China (31571083) to Z.Y., Additional support was provided grants from the Young 1000 Talent Program of China to Z.Y.

---

**S1 Fig.**
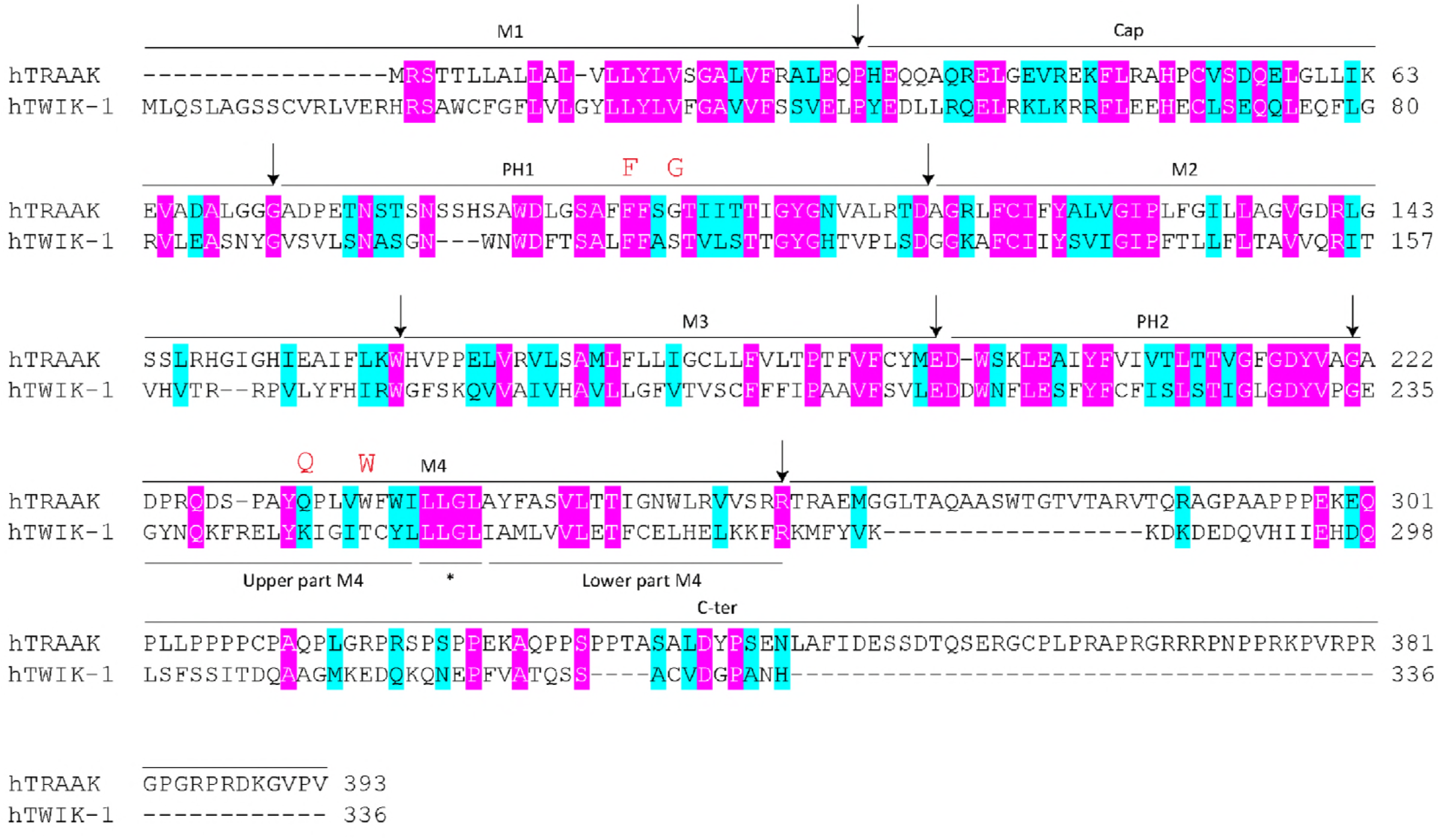
The eight segments of human TRAAK and human TWIK-1. The first transmembrane helix (M1), the two-cap domain (Cap), the first pore helix and selectivity filter (PH1-filter), the second transmembrane helix (M2), the third transmembrane helix (M3), the second pore helix and selectivity filter (PH2-filter), the fourth transmembrane helix (M4) and the C-terminal domain (C-ter) are indicated. F121, G124, Q258 and W262 are highlighted in red.

**S2 Fig.**
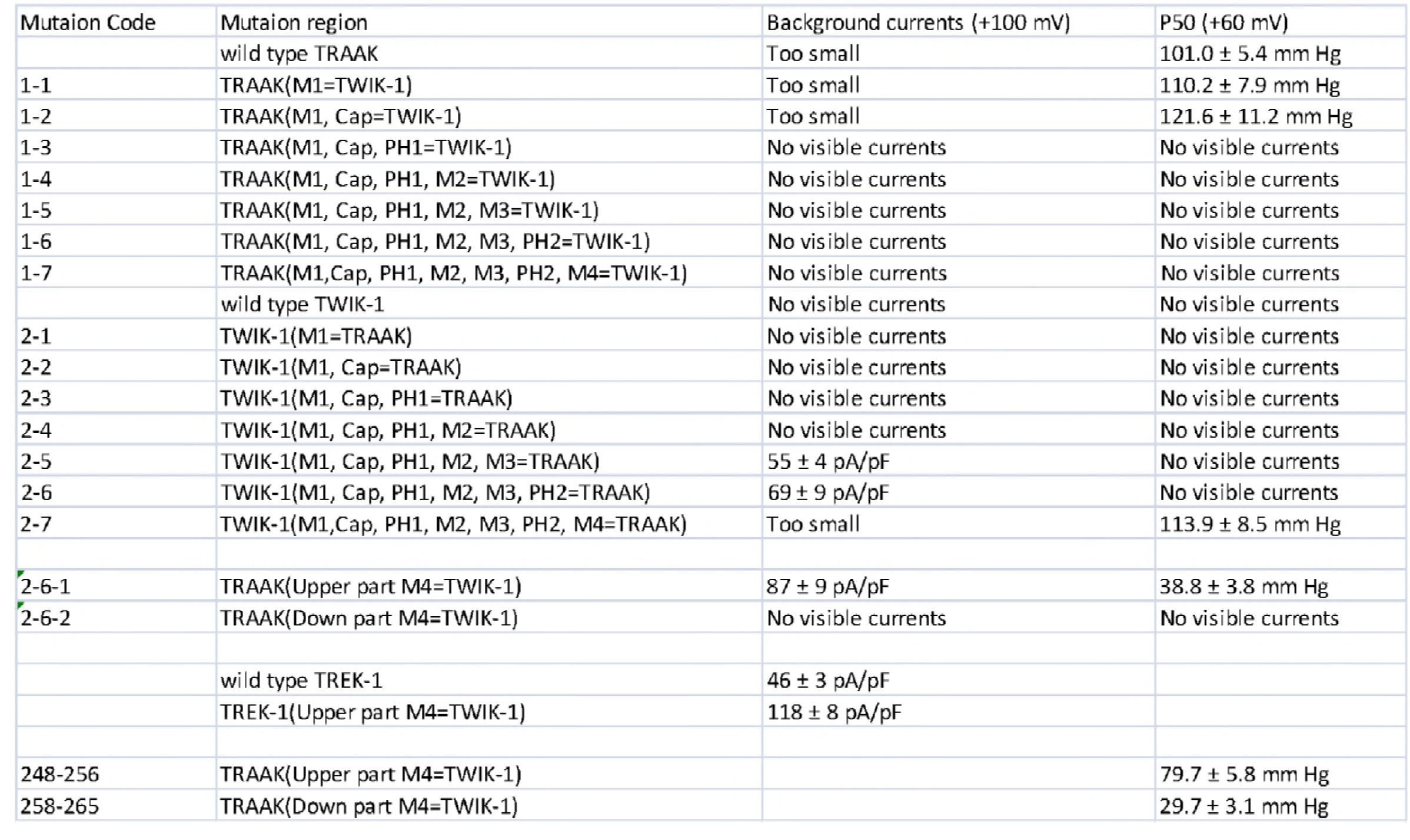
The chimera mutation information and their channel characteristics.

Author Contributions: M.Z. conceived the project and designed experiments. M.Z., C.P. and F.Y performed mutagenesis and electrophysiology experiments. M.Z. and Z.Y. wrote the manuscript.

